# The evolution of the duckweed ionome mirrors losses in structural complexity

**DOI:** 10.1101/2023.09.22.558936

**Authors:** Kellie E Smith, Min Zhou, Paulina Flis, Dylan Jones, Anthony Bishopp, Levi Yant

**Affiliations:** School of Life Sciences, University of Nottingham, Nottingham NG7 2RD, UK; School of Life Sciences, Chongqing University, Chongqing 401331, China; School of Biosciences, University of Nottingham, Sutton Bonington LE12 5RD, UK

**Author notes:** Kellie Smith and Min Zhou contributed equally.

**Keywords:** vestigiality, duckweed, ionomics, evolution, ICP-MS, *Spirodela*, *Landoltia*, *Lemna*, *Wolffiella*, *Wolffia*

## Abstract

**Background and Aims:** The duckweeds consist of 36 species exhibiting impressive phenotypic variation, including the progressive evolutionary loss of a fundamental plant organ, the root. Loss of roots and reduction of vascular tissues in recently derived taxa occur in concert with genome expansions of up to 14-fold. Given the paired loss of roots and reduction in structural complexity in derived taxa, we focus on the evolution of the ionome (whole-plant elemental contents) in the context of these fundamental body plan changes. We expect that progressive vestigiality and eventual loss of roots may have both adaptive and maladaptive consequences which are hitherto unknown.

**Methods:** We quantify the ionomes of 34 accessions in 21 species across all duckweed genera, spanning 70 million years in this rapid cycling plant (doubling times are as low as 24 hours). We relate both micro– and macroevolutionary ionome contrasts to body plan remodelling and show nimble microevolutionary shifts in elemental accumulation and exclusion in novel accessions.

**Key Results:** We observe a robust directional trend in calcium and magnesium levels decreasing from the ancestral representative *Spirodela* genus towards the derived rootless *Wolffia*, with the latter also accumulating cadmium. We also identify abundant within-species variation and hyperaccumulators of specific elements, with this extensive variation at the fine– as opposed to broad-scale.

**Conclusions:** These data underscore the impact of root loss, and reveal the very fine scale of microevolutionary variation in hyperaccumulation and exclusion of a wide range of elements. Broadly, they may point to trade-offs not well recognized in ionomes.

## INTRODUCTION

The duckweeds consist of 36 species exhibiting broad variation, including in recently derived species the progressive evolutionary loss of a fundamental plant organ, the root. This progressive loss of roots is accompanied by an overall reduction in vascular tissues in derived taxa. Given the paired loss of roots and reduction in structural complexity, we here focus on the evolution of the ionome and place it in the context of these fundamental body plan changes.

Consisting of five genera progressively differing in root number and vascular complexity, the duckweeds present broad variation in highly simplified body plans (Figure 1). The earliest diverged lineages*, Spirodela* and *Landoltia* (Fig. 1, top), were originally both considered Spirodela, but are now recognized as distinct (Les & Crawford, 1999; Les *et al*., 2001; Bog *et al*., 2015). The three more recently diverged genera, *Lemna*, *Wolffiella* and *Wolffia*, represent novel forms, with progressively diminished roots and reduced vascular tissues (called nerves) - or none at all (Fig. 1, bottom; Appenroth *et al*., 2013; Tippery *et al*., 2015). The divergence time between rooted *Spirodela polyrhiza* and rootless *Wolffia australiana* is estimated at 70 million years (Park *et al*., 2021). Since this divergence, at least 36 duckweed species have formed (Bog *et al*., 2020; Cao *et al*., 2020), which vary 14-fold in genome sizes. The smallest is an Arabidopsis-scale 158 Mb genome in *Spirodela polyrhiza* (Wang *et al*., 2011; An *et al*., 2018), with the largest genomes in the derived *Wolffia*, exhibiting a radically simplified body plan and diminished vasculature, and no roots (Fig. 1 bottom row; Park *et al*., 2021; Yang *et al*., 2021).

**Figure 1.**
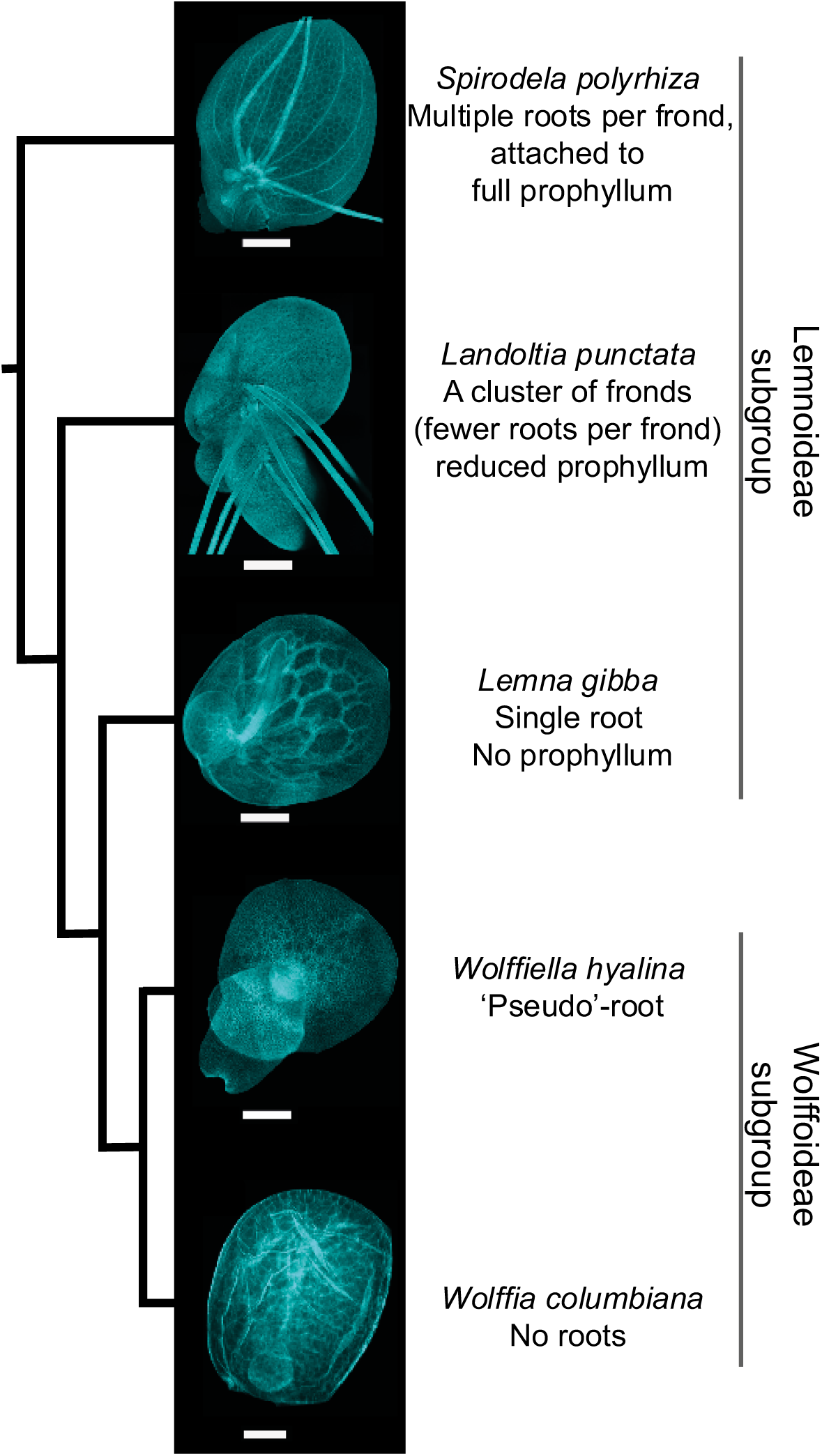
Trajectory from ancestral root-harbouring duckweeds, via vestigiality, to root loss. Ancestral form (above) represented by Lemnoideae: *Spirodela, Landoltia, Lemna*. Derived from (below) shown in Wolffoideae subgroup genera *Wolffiella* and *Wolffia*. All samples were cleared, and stained with Fluorescent Brightener 28 (calcofluor) following the protocol described by (Kurihara *et al*., 2015) and imaged on a Leica TCS SP5 confocal microscope. Scale bars: *Spirodela*, *Landoltia* = 1000 µm; *Lemna, Wolffiella* = 500 µm; *Wolffia* = 100 µm. Cladogram schematic topology based on Tippery et al 2015.

In contrast to vascular land plants, duckweeds have miniscule bodies in direct contact with water and limited to non-existent root systems. This results in small distances for ion translocation (Zhang *et al*., 2009). However, the relative differences in translocation distance can be large: frond sizes of *Spirodela* are >1 cm, but only <1 mm in *Wolffia*. Indeed, in the highly derived Wolffioideae the shrinking of body size and root loss have evolved to maximize growth rate, improve mobility, and enhance adaptability to changing environments (Wang *et al*., 2010; Appenroth *et al*., 2013). We expect that duckweeds, representing this unique example of progressive root reduction through to complete loss, would illustrate a gradient of phenotypic changes resulting in altered internal macronutrient and trace element compositions (Fang *et al*., 2023; Ware *et al*., 2023).

At the fine scale, duckweed habitats differ in their availability of elements; thus adaptation of accessions to their environments can occur through different elemental storage and exclusion strategies (Mkandawire & Dudel, 2007; Zhang *et al*., 2009; Van Dam *et al*., 2010; Lahive *et al*., 2011). Indeed, duckweed tolerance to elemental extremes is an important trait driving adaptive – and sometimes strongly invasive – strategies in the wild (Wang, 1991; Naumann *et al*., 2007; Ekperusi *et al*., 2019). To date, however, the tolerance of only a handful of duckweed accessions to external elemental concentrations has been assessed, with reports focusing on growth vigour vis-à-vis single elements in *Lemna* and *Landoltia* species. Studies quantifying elemental composition are rare, with the broadest study looking at only a single genus, *Wolffia*, with 11 species assessed (Appenroth, 2018). We collected existing reports of duckweed elemental variation; however, serious confounding factors plague interpretation of different studies, due to discordant methods and quantification (Table 1).

**Table 1.** Elemental tissue concentration of duckweeds gathered from the literature. Elements are ordered by type (macro, micro, trace element and heavy metals) reported from the literature and included in our experiment. 1. Appenroth *et al*., 2018. 2. Mkandawire & Dudel, 2007. 3. Van Steveninck *et al*., 1992. 4. Khellaf & Zerdaoui, 2009. 5. Lahive *et al*., 2011. 6. Liu *et al*., 2017. 7. Prasad *et al*., 2001. 8. Leblebici *et al*., 2010. 9. Landolt & Kandeler, 1987.

| Element | Species | Fold variation<br>(literature) | Fold variation (this<br>study, 21 species) |
| --- | --- | --- | --- |
| P | *Wolffia* spp. | 1.7<sup>1,2</sup> | 2.4 |
| K | *Lemna* spp., *Wolffia* spp. | 2.4<sup>1,2</sup> | 3.3 |
| Ca | *Lemna* spp., *Wolffia* spp. | 3.3<sup>1,2</sup> | 11.4 |
| Mg | *Lemna* spp., *Wolffia* spp. | 3.1<sup>1,2</sup> | 19.5 |
| Na | *Lemna* spp., *Wolffia* spp. | 29.5<sup>1,2</sup> | 27.4 |
| Fe | *Lemna* spp., *Wolffia* spp. | 21.8<sup>1,2</sup> | 111.0 |
| Zn | *Lemna gibba*, *Lemna minor*,<br>*Landoltia punctata*, *Wolffia* spp. | 87.3<sup>1,3,4,5</sup> | 149.6 |
| Mn | *Spirodela polyrhiza*, *Wolffia* spp. | 27.3<sup>1,6</sup> | 4.5 |
| Cu | *Lemna trisulca*, *Lemna gibba*,<br>*Lemna minor*, *Wolffia* spp. | 15.7<sup>1,7,8,9</sup> | 7.6 |
| Cd | *Landoltia punctata* 6001, *Lemna*<br>*minor*, *Lemna gibba*, *Spirodela*<br>*polyrhiza* sp., *Wolffia globosa*. | 5,900<br><sup>1,10,11,12,13,14</sup> | 27.3 |

Here we bridge this gap, reporting whole-plant ionome compositions in 34 duckweed accessions spanning 21 species and representing the worldwide range of all five duckweed genera (Figure 2; Supplementary Table 1). We place these data into an evolutionary context, focusing on 11 key macro-, micro-, and trace elements, contrasting microevolutionary variation (accession level, within species variation) with macroevolutionary (between genera) trends. These results reveal extensive ionomic variation at both the within-species and between-genus levels, with particularly clear trends for Ca and Mg accumulation differences, as well as possible excess Cd accumulation in the rootless *Wolffia/Wolffiella*. We discern a broad evolutionary trajectory toward very low levels of essential Ca and Mg—as well as increased Cd accumulation—in the recently derived rootless species. This suggests a potentially deleterious consequence associated with the root loss and body-wide vasculature reduction.

**Figure 2.**
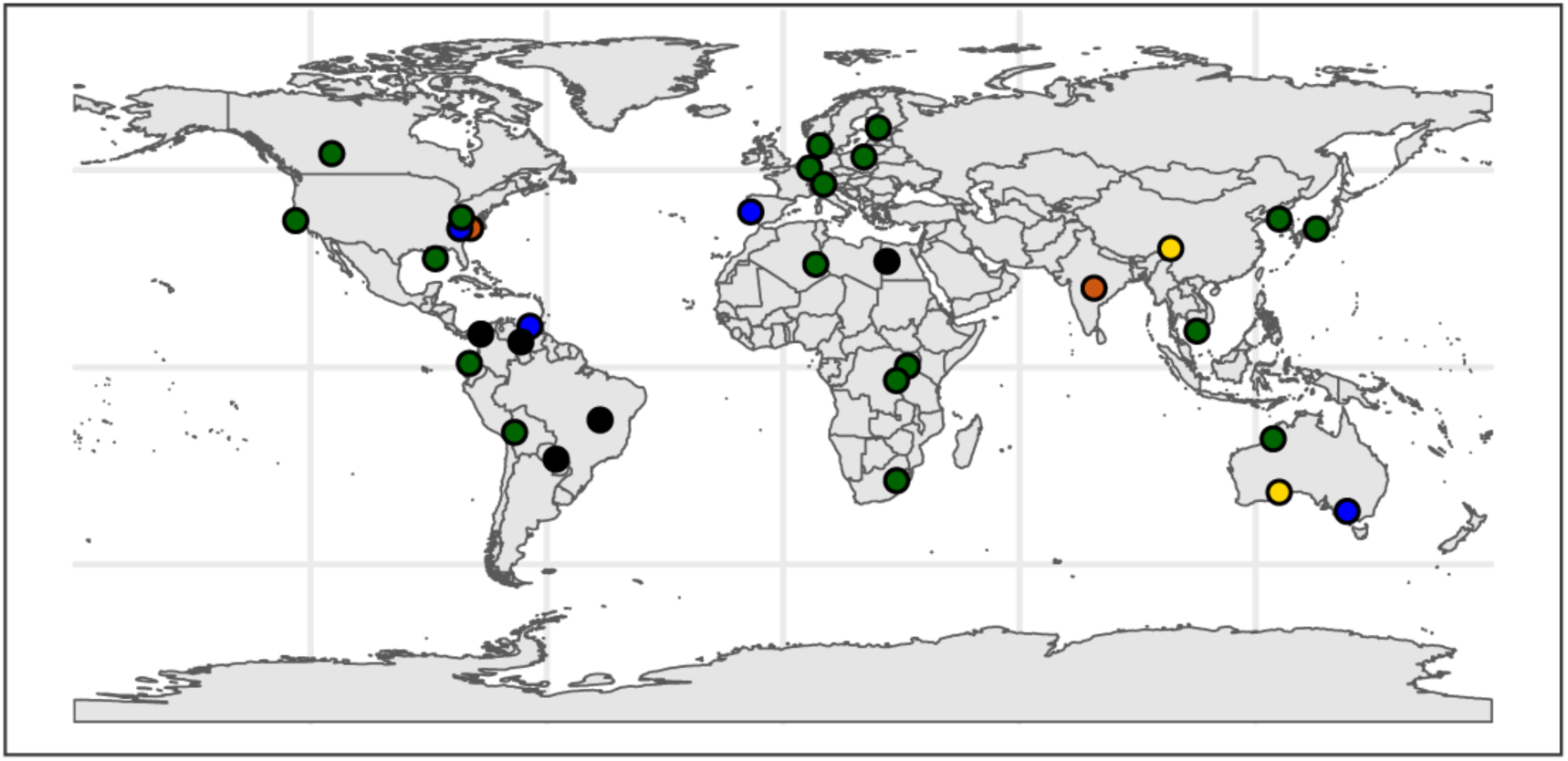
Sampling of worldwide duckweeds for ionomic panel. Dots indicate sample origin locations: *Lemna*=green, *Landoltia*=yellow, *Spirodela*=black, *Wolffiella*=orange, *Wolffia*=blue.

## MATERIALS AND METHODS

### Plant Growth and Care

Duckweed accessions were grown from single isolates or 5-10 individuals, depending on size of duckweeds, in 100 ml nutrient media (N media) in individual sealed sterile glass conical flasks. N media as described in Appenroth *et al*, 1996 (KH_2_PO_4_ (0.15 mM), Ca(NO_3_)_2_ (1 mM), KNO_3_ (8 mM), MgSO_4_ (1 mM), H_3_BO_3_ (5 µM), MnCl2 (13 µM), Na_2_MoO_4_ (0.4 µM), FeEDTA (25 µM). Element concentrations of supplied N media including presence of other trace elements were measured by ICP-MS and is presented in Dataset S1. Weekly media changes were performed, with rinses in MQ water to regulate nutrient composition availability. Plants were grown at 100 µmol m^-2^ s^-1^ at 22°C/18°C with a 16-hour day/night cycle. Four-week-old duckweed cultures were washed on plastic sieves using a three-step protocol two minutes each of MQ water, CaCl2 and Na-EDTA and harvested into individual samples from flasks of individual populations. These were harvested for ICP-MS analysis on day one, three and five after media change, n=6 per time point. Four-week-old cultures are clonally reproduced and therefore suitable replicates, given the very low generational variation and low mutation rates shown in duckweed mutation accumulation experiments (Xu *et al*., 2019).

### Imaging and Microscopy

All samples were cleared and then stained with Fluorescent Brightener 28 (calcofluor) following the protocol described by (Kurihara et al., 2015) and imaged on a Leica TCS SP5 confocal microscope. In short, plants were cleared, based on the ClearSee procedure described by Kurihara et al. (2015), modified slightly. As fluorescent markers were not being used, plants were fixed overnight in ethanol and acetic acid (3:1 v/v) rather than paraformaldehyde, as this reduced the toxicity and requirement for vacuum infiltration, which can be damaging to the air spaces. Plants were then rinsed three times in RO water and left for 30 min, then RO water was replaced with ClearSee solution (10% Xylitol, 15% Sodium Deoxycholate, 25% Urea; Kurihara et al., 2015) and left to clear for 2 weeks. Prior to imaging, plants were stained for 1 h with calcofluor in ClearSee (100 μg/ml), and then washed in ClearSee for 1 h. Imaging was carried using a confocal laser scanning microscope (Leica SP5), using a 405 nm diode laser at 12% and hybrid detector with a range of 440–450 nm, gain of 25%, and pinhole of 0.5 AU.

### Quantification of elemental tissue concentrations

For ICP-MS we used a method adapted from Danku et al., 2013. Briefly, 5-20 mg (fresh weight) was harvested per sample, placed in Pyrex test tubes and dried at 88°C for 24h. Then the dry weight was recorded and 1 ml concentrated trace metal grade nitric acid Primar Plus (Fisher Chemicals) spiked with an internal standard was added to the samples that were further digested in DigiPREP MS dry block heaters (SCP Science; QMX Laboratories) for 4h at 115°C. Prior to the digestion, 20 µg/ L of Indium (In) was added to the nitric acid as an internal standard for assessing errors in dilution, variations in sample introduction and plasma stability in the ICP-MS instrument. Then 0.5 ml of hydrogen peroxide (Primar, for trace metal analysis, Fisher Chemicals) was added to the samples and they were digested for additional 1.5 hr at 115°C. After digestion, samples and blanks were diluted to 10 ml with Milli-Q Direct water and elemental analysis was performed using an ICP-MS, PerkinElmer NexION 2000 with 22 elements monitored (Li, B, Na, Mg, P, S, K, Ca, Cr, Mn, Fe, Co, Ni, Cu, Zn, As, Se, Rb, Sr, Mo, Cd and Pb) in the collision mode (He). To correct for variation between and within ICP-MS analysis runs, liquid reference material was prepared using pooled digested samples, and run after every nine samples in all ICP-MS sample sets. The calibration standards were prepared from single element standards solutions (Inorganic Ventures; Essex Scientific Laboratory Supplies Ltd, Essex, UK). Sample concentrations were calculated using external calibration method within the instrument software. Further data processing including calculation of final elements concentrations (in mg/kg) was performed in Microsoft Excel. Log transformations, Z-score calculations and graphical representation was performed using R (version 3.0.2 “Frisbee Sailing”; R Development Core Team, 2013; see http://www.R-project.org) and R Studio (v 1.0.136) was used for all statistical analyses.

## RESULTS

### Broad scale evolution of the ionome

We focus on ionomes from day five after media change (Fig. 3), which is representative of other time points (None of the 11 elements upon which we focus were significantly different across days by ANOVA). The full data set is given in Dataset S2; elements we considered for further analysis shown in Supplementary Figure 1. Concentrations were consistent for all elements for all accessions between time points except for a handful of elements in certain accessions depicted in Supplementary Figure 1. These exceptions show a small minority of accessions decreasing in K, Ca, Fe and Cd and others still increasing *e.g* Ca, Cu, Fe. For accumulators showing the latter pattern, for example *Sp. intermedia* 9227, maximum concentration capacity of Ca on day one after media changes was still not reached, despite high nutrient provision throughout a four-week experimental period, and the accession could still prolong uptake.

**Figure 3.**
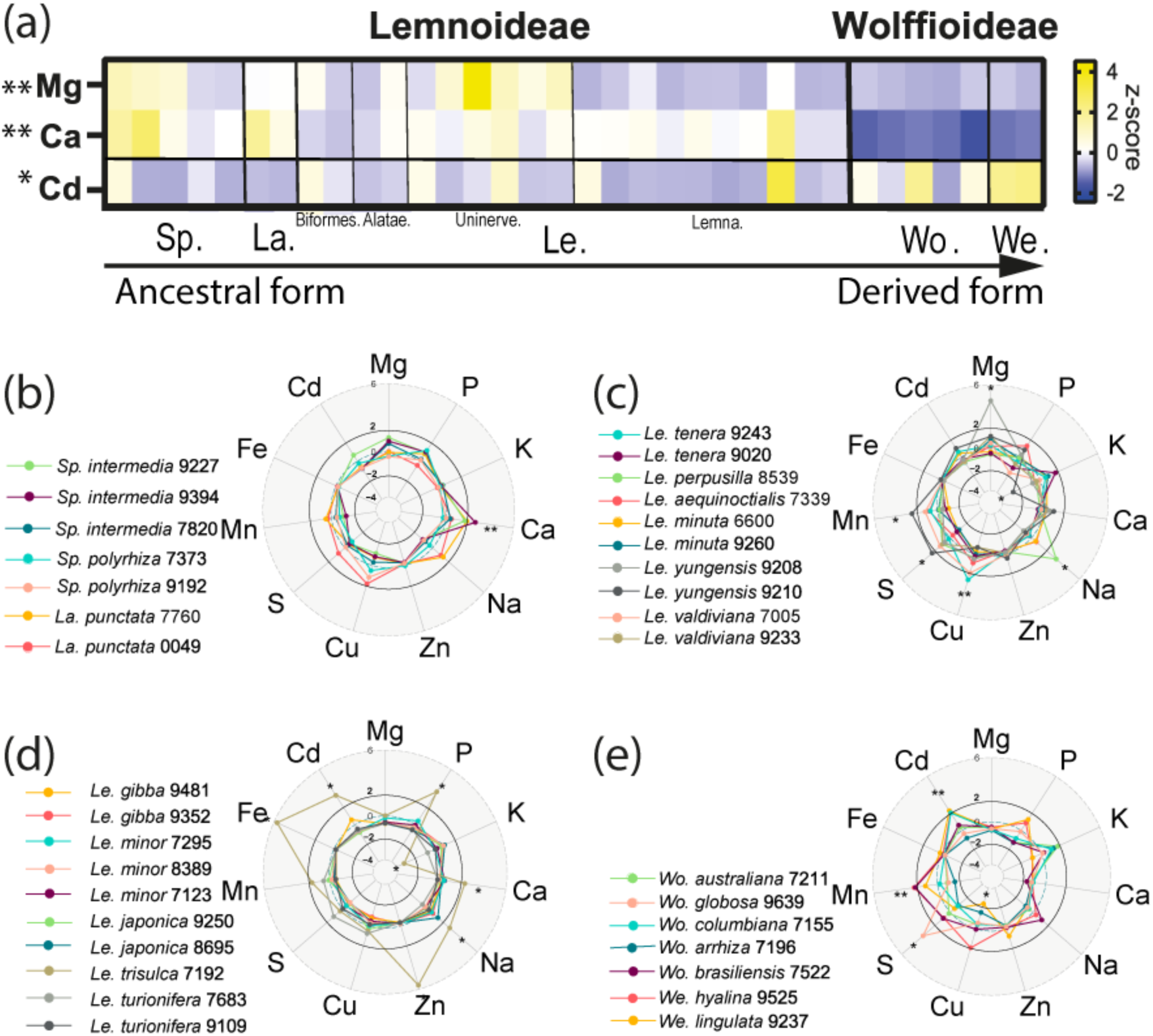
The evolution of the duckweed ionome across genera, species, and accessions. (a) Relative levels of elemental accumulation across rootless and rooted subgroups, respectively. The heat map is coloured by z-scores for the four most differentially accumulated elements by significant ANOVA Tukey ** significant *p*<0.01 and * *p*<0.05 ANOVA with post-hoc Tukey test differences between Wolffioideae and Lemnoideae. Z-scores (number of standard deviations away from the mean) were generated for each element using log10 transformation of mg/kg values on day 5. X axis i s arranged with basal forms on the left and derived on the right. Separating lines indicate genus and subgroup boundaries. We. = *Wolffiella* (2), Wo. = *Wolffia* (5) Le. = *Lemna* (20), La. = *Landoltia* (2) and Sp. = *Spirodela* (5). Within Lemna subgroups Biformes, Alatae, Uninerve and Lemna marked left to right; **(b-e)** Radar plots showing differences in ionome profiles between individual accessions. **(b)** *Spirodela* and *Landoltia*; **(c)** *Lemna* Biformes, Alatae and Uninerves; **(d)** *Lemna* sect. Lemna; **(e)** *Wolffiella* and *Wolffia* species. Species are ordered in the panels according to (Tippery *et al*., 2015) with most ancestral representative at top left through to most derived in bottom right. Numbers after species represent clone numbers. Asterisks represent significant increase or decrease +/− 2 relative to all normalised element concentrations for all species based on mean and SD. The complete data set of 17 elements and three timepoints is given in Dataset S2.

In the overall dataset of 34 accessions, the broadest contrast observed was between the Lemnoideae and Wolffioideae (rooted and rootless, respectively) for Ca, Mg and Cd accumulation (Figure 3a). All ancestral representatives (rooted) Lemnoideae (*Spirodela*, *Landoltia* and *Lemna*) consistently exhibit 2-3x higher Ca content relative to the derived rootless Wolffioideae (p =<0.01; log10, ANOVA with post-hoc Tukey test). Similarly, on average, Mg accumulation was 1.8x higher in the rooted species relative to the rootless *Wolffia* and *Wolffiella*. Further variation was within the *Lemna* genus, with the highest levels of Mg in the uninerve subgroup of *Lemna* (Fig 3a, Fig 4), which includes the invasive *Lemna minuta,* and *Lemna yungensis* species, as described in (Tippery *et al*., 2015 and Bog *et al*., 2020). This association of Mg accumulation with increased root vasculature (as well as frond vasculature in *Lemna*) stands in strong contrast to the uniformly very low Mg in rootless Wolffioideae.

**Figure 4.**
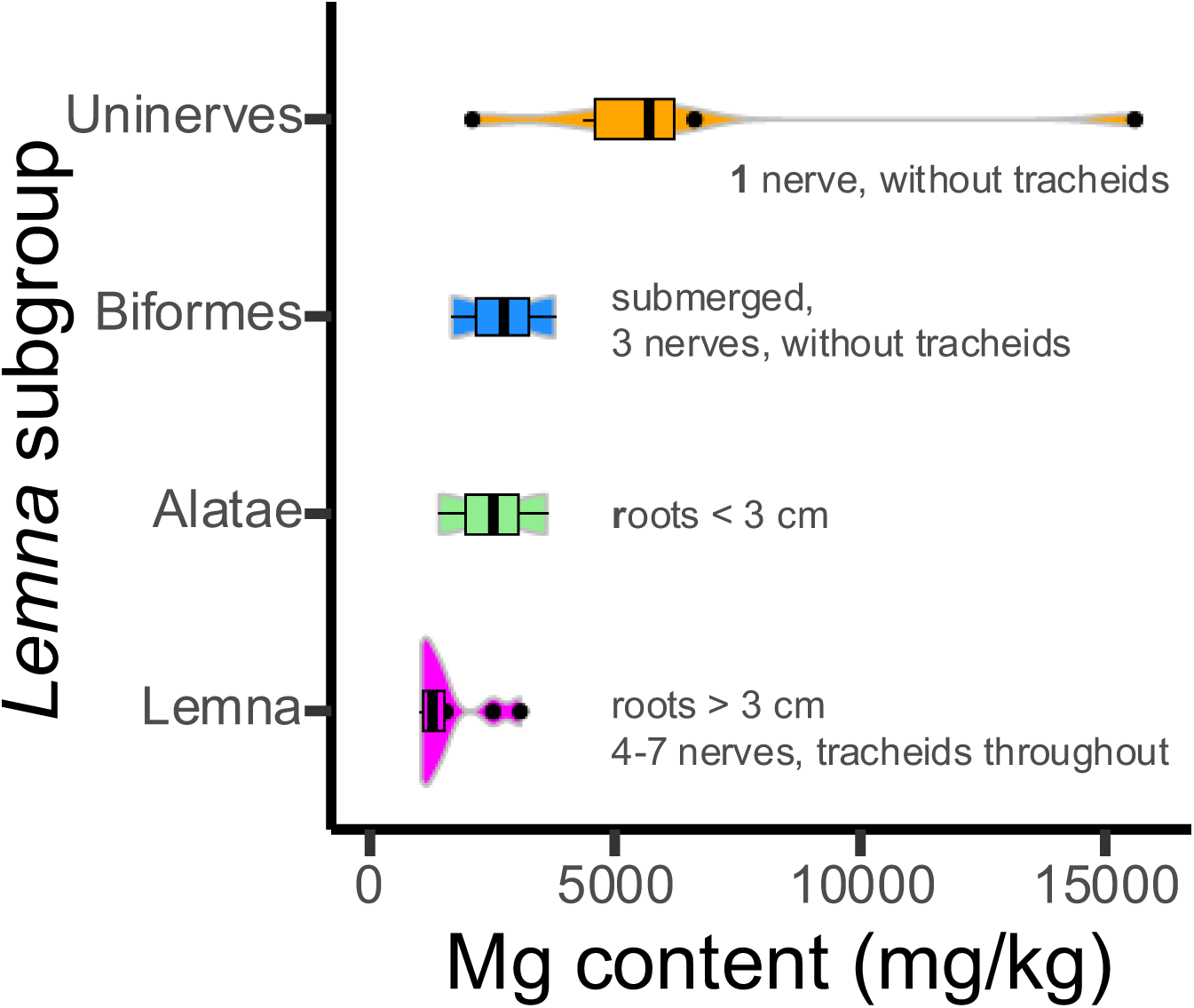
Increased Mg content mirrors the reduction of frond vasculature within *Lemna*. Four subgroups of *Lemna* representing highest Mg content in the species with most reduced vasculature at sect Uninerves, with transition subgroups Biformes and Alatae and most developed frond vasculature in sect Lemna, with reduced Mg. Mg content is plotted from day five averaged values for each accession within each group: Uninerves *n*=6, Biformes *n*=2, Alatae *n*=2, Lemna *n*=10. Groups ordered and described according to (Landolt, 1986; Tippery *et al*., 2015). Violin plots represent spread of data for each group with middle line plotting mean.

Interestingly, the *Lemna* sect of *Lemna*, that is more closely related to rootless duckweeds, do have similar Mg accumulation to the Wolffioideae species, indicating a gradient of differential Mg accumulation in the *Lemna* genus (Fig 1, Fig 3 a, d). This gradient does not necessarily correlate with strict overall inferred ancestral and derived forms (Wang *et al*., 2011; Tippery *et al*., 2015) and root vascular complexity is not sufficiently varied between rooted duckweeds to account for this (Ware *et al*., 2023). Instead, specific Mg uptake in the uninerve clade of Lemna may be associated with their reduced frond vascular complexity (Fig. 3a, Fig 4). With typical frond nerves up to 16 in number in *Spirodela* and between 3-7 in other *Lemna* species (Les *et al*., 2002), only one nerve is present in *L. yungensis* and *L. minuta*, with *L. yungensis* having the longer nerve of the two, covering almost the entire distance from the node to apex of its frond (Landolt, 1980; Crawford *et al*., 1996). Vein leaf density is a factor involved in phloem transport capacity of nutrients (Zhang *et al*., 2015), indicating that uninerve *Lemna* may have reduced capacity to transport elements across the frond, compared to other *Lemna*, and possibly results in bottlenecks in other organs, *e.g* the root. Cadmium concentrations varied significantly between rooted and non-rooted duckweeds (Fig. 3a; p < 0.05; log 10, ANOVA with post-hoc Tukey test) in a manner inverse to Ca and Mg. The unrooted Wolffioideae species (especially *Wolffiella*) showed the highest Cd concentrations (Fig. 3). Only the submerged *Lemna trisulca* exhibited comparably high Cd to the Wolffioideae (Fig. 3).

While some variation in mineral content among *Wolffia* species has been reported by Appenroth *et al*, (2018), *Wolffiella* have received little attention and can be under-reported due to clones not being readily available (Landolt, 1986; Cao *et al*., 2020). Rootless species exhibiting variation in at least two elements included *Wolffiella lingulata, Wolffiella hyaline,* and *Wolffia brasiliensis* (Fig. 3e). In contrast, the species in our panel from the multi-rooted more ancestral duckweed representatives *Spirodela* and *Landoltia* showed the greatest ionomic consistency across all accessions (Fig. 3b). *Spirodela* species had the highest Ca tissue content in our panel, but other elements were not as variable between accessions. As Ca was kept sufficiently available in our experiment through the media refreshes, it is likely that rooted duckweeds use their roots as a storage compartment when such nutrients are replete, and as a result accumulate more when compared with their rootless counterparts.

### Fine scale ionome variation and identification of extreme accumulators in Lemna

We observe the greatest within-genus ionome variation in the *Lemna* genus (*n*=20 accessions, 6 biological replicates of each; Fig. 3c-d). *Lemna* also harbours several extreme accumulators, each standing as outliers for the accumulation of three or more elements. *Lemna trisulca* 7192 has a submerged growth pattern and accumulated the greatest number of elements in amount and number from the panel, showing very high tissue concentrations of four essential elements: P, Ca, Zn, Fe as well as Cd, and low K levels (Fig. 3d). *Lemna yungensis* 9210 accumulated high S and Mn, and also exhibited low K (Fig. 3c). Indeed, K levels trend negatively against the enhanced accumulation of other macro– and microelements in both *L. trisulca* and *L. yungensis* (Table 2; Supplementary Figure 4). Both *L. trisulca* and *L. yungensis* accumulated over 1000 mg/kg dry weight for several elements and can be considered to be hyperaccumulators (Zayed *et al*., 1998; Zhang *et al*., 2009). This suggests that these two species may have potential to be used in combination to alleviate multi-elemental toxicity in watercourses. *Lemna trisulca* accumulated greater Zn and Cd than floating species, possibly because of increased absorption through submerged fronds. Although *L. trisulca* had the greatest variation and maximal micronutrient levels, the associated high Cd accumulation may be problematic for any applications in nutrition. It is also unclear whether this trait is common in other *L. trisulca* accessions due to limited availability of clones in stock centres.

**Table 2.** Element pairs significantly correlated across 34 duckweeds at three time points. R^2^ values correspond to positive or negative Pearson correlations.

| Element | $R^2$ |
| --- | --- |
| Fe/K | -0.58 |
| Zn/K | -0.525 |
| P/K | -0.443 |
| Mn/K | -0.346 |
| Fe/Mn | 0.355 |
| Mg/Ca | 0.349 |
| Zn/Mn | 0.339 |

At the within-species level we detected variation at the level of several accession pairs, most obviously between *L. yungensis* accessions (Fig. 3c). Notably, *L. yungensis* 9208 greatly accumulated Mg, and *L. yungensis* 9210 exhibited extreme accumulation of S and Mn, but low K. Given these accessions are closely related and were both originally isolated from the same region in Bolivia, one might expect more similar ionome profiles, but instead our data show that duckweeds exhibit strongly contrasting local variation in elemental uptake. Interestingly, this region of Bolivia is reported as atypically harsh for duckweed, growing on sheer rock faces with waterfall spray with low nutrient availability (Landolt, 1998). It will be valuable to characterise *L. yungensis* species further, in order to determine the genetic basis for their adaptation to specialised habitats.

## DISCUSSION

The broad variation we observed in duckweed ionomes at levels of genera, species, and sister accessions is presumably in large part due to both morphological differences and adaptation to micro-environments. The most robust differences were at the genus level for Ca, Mg and Cd. The accumulation difference for Ca is perhaps explained in part by the storage mechanism of Ca, as Ca oxalate (CaOx) within frond crystal ultrastructures in rooted genera, in the fronds of *Spirodela* and *Lemna*, (Landolt & Kandeler, 1987) and also in the root of *L. minor* (Franceschi, 1989, Mazen *et al*., 2003). In contrast, Wolffioideae species have soluble Ca in cell sap and accordingly also cannot store excess Ca in roots (Landolt & Kandeler, 1987; Appenroth *et al*., 2017); thus Ca and Mg may be lower in Wolffiodieae as they lack roots as a storage organ.

Given the broad contrasts in Ca between genera, it is interesting to consider these results alongside the importance of roots for elemental uptake and segregation of individual elements between frond and roots in duckweed species. The excision of roots makes only a modest change to the frond ionome, showing roots are vestigial and overall not required for nutrient uptake in replete media conditions (Ware *et al*., 2023). Whilst removal of roots surprisingly increased elemental composition in some cases, the picture is more complicated as rootless species do not naturally exhibit elevated Mg or Ca in our data, indicating evolutionary adjustment of ion homeostasis upon root loss. Indeed, the *Wolffia* genome harbours a derived complement of Ca export and cell wall thickening genes, possibly minimizing potential for apoplastic transport, which coupled with inability for storage as CaOx, results in less specialised mechanisms to manoeuvre and store Ca content overall (Michael *et al*., 2020). In contrast, the *Lemna* species *Le. aequinoctialis*, *Le. minuta* and *Le. minor* exhibit marked Ca accumulation (storage) to alleviate Mg toxicity from a contaminated mine and on high Mg:Ca ratio media or wastewater (Van Dam *et al*., 2010; Paolacci *et al*., 2016; Walsh *et al*., 2020). This suggests specific adaptation of Ca storage and transport mechanisms to particular ionomic challenges.

The relative accumulation of Cd, especially in *Wolffiella* species compared to the rooted species, is somewhat surprising, as it would be expected that Cd accumulation would be detrimental to minuscule plants with no root segregation away from photosynthetically active tissue. Here we find increased Zn accumulation associated with Cd and Fe (Fig S4), highlighting complex element interactions, also found in *S. polyrhiza* and sunflower (Lopes Júnior *et al*., 2014; Su *et al*., 2017); however it is unclear whether enhancement of trace metals in *Wollfia* offer any evolutionary advantage. *Wolffia* species also exhibit tolerance to As and have been considered as candidates for phytoremediation, accumulating higher than Lemnoideae duckweeds (Zhang *et al*., 2009). Additionally, *Wolffia* has moderate tolerance to Cd and increased accumulation capacity even in extreme concentrations (>200 µM). In fact, a handful of *Wolffia* species show Cd uptake in as little as 30 minutes from solution via apoplast which increases linearly with Cd concentration (Boonyapookana *et al*., 2002; Xie *et al*., 2013). We therefore speculate that loss of roots could have reduced control of heavy metal uptake but Wolffioideae perhaps evolved higher tolerance mechanisms to its toxicity. *Wolffia* also clearly has the potential for heavy metal accumulation at much greater dosage than shown here, perhaps also through adaptation to contaminated habitats (Zhang *et al*., 2009).

Our data reveal that the genus with the greatest diversity of specific accumulators was *Lemna*. The *Lemna* accessions with most extreme ionomes, *Le. trisulca* 7192 and *Le. yungensis* 2908, also harbour the most divergent root complexity in terms of structure, compared even to other *Lemnas*. *Lemna trisulca* is characterised by a submerged growth habit but smaller cortical cells giving a thin, reduced root compared to other *Lemna* species, and *Le. yungensis* 9208 often displays an additional layer of cortical cells and irregularly large extracellular airspaces in the root cortex (Ware *et al*., 2023). Thus, these differential root vasculature components, coupled with minimal frond vasculature, may play a role in producing the contrasting elemental profiles observed. Differential accumulation experiments between frond and root segregation of elements including Mg in these species would be required to see if there is a bottleneck of storage in the roots, and limited in the frond, due perhaps to reduced vasculature.

A greater appreciation for duckweed variation in micronutrients Ca, Mg, Fe and Zn is clear from our study, with particular accessions acting as hyperaccumulators for multiple nutritionally-relevant elements. This is not the case for trace elements such as Na and Cu (and especially Mn and the heavy metal Cd), where tissue concentration variation was less dramatic than seen in other reports (Table 1). This is likely due to the combined effect of low presence of these elements in our supplied media or that comparisons across literature are confounded by variables disallowing truly quantitative comparisons between studies. This is particularly evident for Cd, which we supply only in trace amounts (Dataset S1), whereas external Cd concentrations vary 500-fold between studies.

Synthetic biology, including the tailoring of ionomic profiles in duckweeds, is an important goal of the duckweed research community (Lam & Michael, 2022). Interestingly, the *Spirodela* genome sizes are the smallest and the ionomes the least variable among all duckweeds here (Wang et al., 2011; An et al., 2018); additionally, the amenability of *Spirodela* to genetic transformation (Yang *et al*., 2018a; Yang *et al*., 2018b) makes it a strong candidate as a minimal scaffold for synthetic biology. We additionally suggest that because their ionomic profiles are so variable, the larger genome-harbouring species will be particularly valuable to mine natural variation to inform transgenic approaches in the smaller, highly tractable *Spirodela* genome.

### Conclusions

Here we detailed broad– and fine-scale diversity for the accumulation of physiologically and nutritionally important elements across all five duckweed genera. This variation is associated with dramatic morphological reductions in fundamental plant organs and genome expansions. Thus, disentangling the concurrent effects of dramatic genome size expansions, organ reduction, and ecological adaptations will be a great challenge. However, at the more microevolutionary scale, within-species, accession-level variation points to clear promise in mapping alleles responsible for this observed variation.

One might speculate that the observed ionomic changes may be a maladaptive spandrel associated with root loss in derived taxa, but it is hard at this point to identify what the exact trade-off may be; this is for dedicated mechanistic and ecological work on the rootless taxa. Beyond highlighting these enigmatic correlates of root loss and the consequences of organ loss and vestigiality, this work serves to establish phenotypic variation across the ionome at both the fine and broad scale. This serves as a basis for future genomic characterisation of causal alleles, as well as rational development of targeted duckweed lines for both important nutritional and phytoremediation goals.

## SUPPLEMENTARY INFORMATION

**Supplementary Figures 1-4:** Available as a separate file

**Supplementary Table 1:** Summary of accessions used in the ionomics panel with corresponding Landolt collection codes and place of origin

**Dataset S1:** Summary elements present in N media, as measured by ICP-MS

**Dataset S2:** All ionomics data (mg/kg) for 22 elements for 34 accessions on day 1, 3, 5 post media change quantified by ICP-MS

## ACKNOWLEDGEMENTS

We thank Todd Michael for helpful comments on an early version of this manuscript. We thank Walter Lämmler for supplying the duckweed used in these experiments. All clones were obtained from the Landolt Duckweed Collection in Zürich, Switzerland. We thank Matt Kent for help with R, Alex Rhodes for his assistance in exploratory analysis, and Alex Ware for cultivating plant material and maintaining the Nottingham Duckweed collection. LY conceived of and oversaw the study, interpreted the data, and secured funding. KES and LY wrote the manuscript with input from all authors. MZ performed the experiments with assistance from AB and PF. KES analysed the data. DHJ performed microscopy. All authors read and contributed to the final manuscript.

## DATA ARCHIVING

The data are given as supplemental information to the article.

