## Supplementary_figures_1_to_4 for "The evolution of the duckweed ionome mirrors losses in structural complexity"

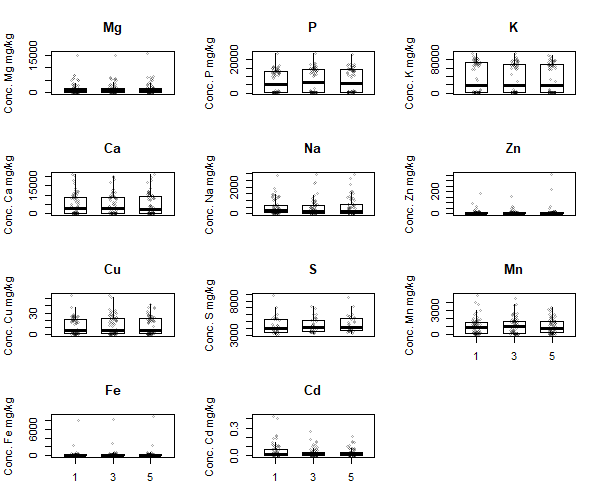

**Supplementary Figure 1.** **Raw elemental composition of duckweed whole plants between day 1, 3 and 5 following media change by ICP-MS.** Boxplots show median value and upper (75%) and lower (25%) quartile of averages from 34 accessions, *n*=6 for each accession at each time point. Eleven elements are shown as independent boxplots, Mg, P, K, Ca, Na, Zn, Cu, S, Mn, Fe and Cd. The following elements were below the limit of quantification (LOQ) and limit of detection (LOD) in duckweed tissue samples using ICP-MS: Selenium LOD = 0.86 ppb/ LOQ = 2.85 ppb, Arsenic LOD = 0.034 ppb/ LOQ = 0.12 ppb. Lithium, Chromium and Lead were not above blank levels in analysed duckweed species.

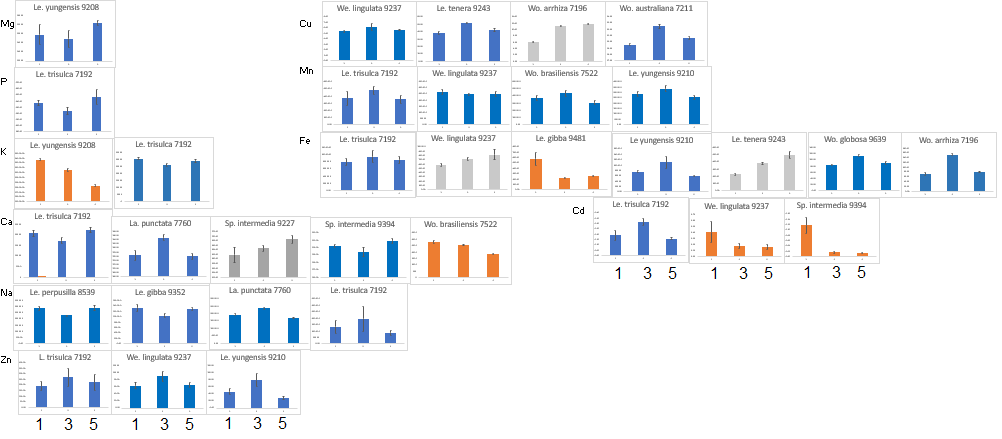

**Supplementary Figure 2 Outlier accessions with dynamic elemental concentrations over sampling days 1, 3, and 5 after media change.** Bar plots in blue show overall stable concentrations, those in grey show an increase over time and those in orange show a decrease over time. Error bars indicate standard error from 6 biological replicates from separate populations.

A. B.

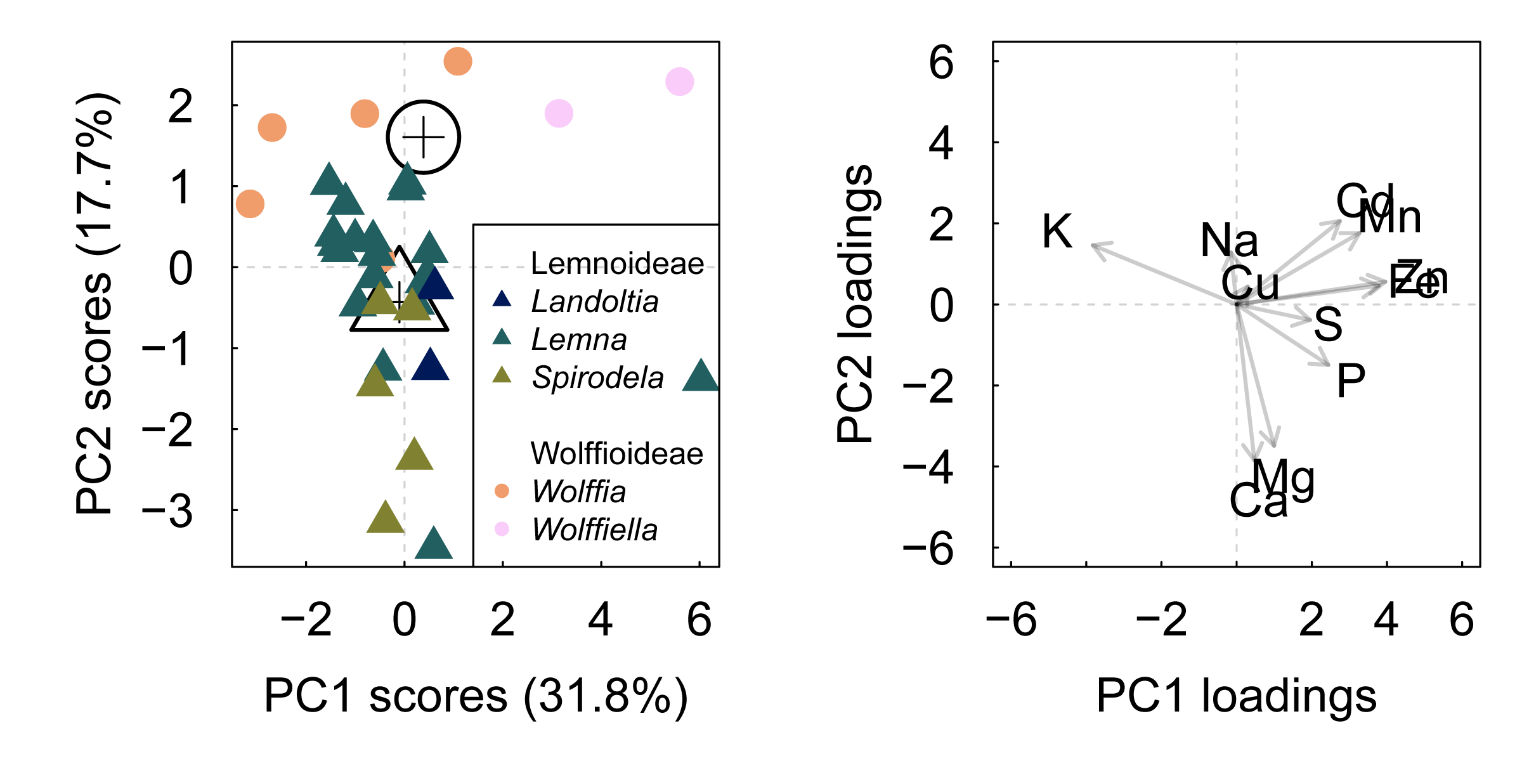

**Fig. S3 Principal component analysis for 11 plant macro- micronutrients and heavy metals.**

**A.** Variable plot explaining 49.5% of the variation in PC1 and PC2 using 11 elemental concentrations averaged from three time points and colored by genus. Genera are further grouped into rooted (Lemnoideae) as triangles and rootless subgroups (Wolffiodeae) as circles. Centroids are generated from means of rooted and rootless subgroups**. B.** PCA plot for 11 elemental tissue concentrations averaged from three days (1,3,5) for 34 accessions. *L.trisulca* is the major outlier across species and has been removed from this dataset to reveal underlying variation across elements and genera.

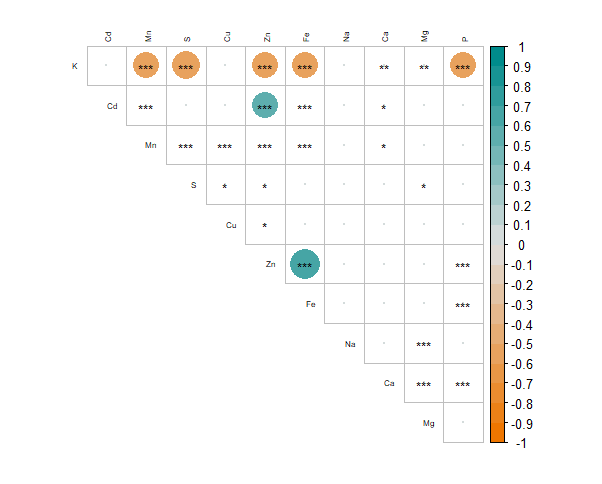

**Supplementary Figure 4 Intensity and direction of correlations between 11 elements in 34 duckweed species.** Significant correlations of elements by Pearson correlation marked as *** for p values <0.001, ** for <0.01 and * for <0.05. Positive relationships indicated by large blue circles r^2^ values >0.3 and negative by orange circles with r^2^ values <-0.3.
