## Supplementary_table for "The evolution of the duckweed ionome mirrors losses in structural complexity"

**Supplementary Table 1.** Accessions studied in this work.

| **Species** | **Landolt Code** | **Location** |
| --- | --- | --- |
| *Wolffiella lingulata* 9237 | WL9237 | Africa |
| *Wolffiella hyalina* 9525 | WA9525 | India |
| *Wolffia brasiliensis* 7522 | WB7522 | North Carolina, USA |
| *Wolffia arrhiza* 7196 | WA7196 | Portugal |
| *Wolffia columbiana* 7155 | WC7155 | Florida, USA |
| *Wolffia globosa* 9639 | WG9639 | Venezuela |
| *Wolffia australiana* 7211 | WA7211 | Victoria, Australia |
| *Lemna turionifera* 9109 | LT9109 | Podlaskie, Poland |
| *Lemna turionifera* 7683 | LT7683 | Kyonggi-Do, Korea |
| *Lemna trisulca* 7192 | LT7192 | Kigezi, Uganda |
| *Lemna japonica* 8695 | LJ8695 | Kyoto, Japan |
| *Lemna japonica* 9250 | LJ9250 | Etela-Suomi, Finland |
| *Lemna minor* 7123 | LM7123 | Saskatchewan, Canada |
| *Lemna minor* 8389 | LM8389 | Transvaal, South Africa |
| *Lemna minor* 7295 | LM7295 | Libya |
| *Lemna gibba* 9352 | LG9352 | Hessen, Germany |
| *Lemna gibba* 9481 | LJ9481 | Denmark |
| *Lemna valdiviana* 9233 | LV9233 | Esmeralda, Ecuador |
| *Lemna valdiviana* 7005 | LV7005 | Florida, USA |
| *Lemna yungensis* 9210 | LY9210 | La Paz, Bolivia |
| *Lemna yungensis* 9208 | LY9208 | La Paz, Bolivia |
| *Lemna minuta* 9260 | LM9260 | Trentino Alto Adige, Italy |
| *Lemna minuta* 6600 | LM6600 | California, USA |
| *Lemna aequinoctialis* 7339 | LA7339 | Burundi |
| *Lemna perpusilla* 8539 | LP8539 | Virginia, USA |
| *Lemna tenera* 9020 | LT9020 | Northern Territory, Australia |
| *Lemna tenera* 9243 | LT9243 | Ca Mao, Vietnam |
| *Landoltia punctata* 0049 | LP0049 | Sichuan, China |
| *Landoltia punctata* 7760 | LP7760 | South Australia |
| *Spirodela polyrhiza* 9192 | SP9192 | Cordoba, Columbia |
| *Spirodela polyrhiza* 7373 | SP7373 | Egypt |
| *Spirodela intermedia* 7820 | SI7820 | Formosa, Paraguay |
| *Spirodela intermedia* 9394 | SI9394 | Sucre, Venezuela |
| *Spirodela intermedia* 9227 | SI9227 | Bahia, Brazil |
